# ESM-PVP: Identification and classification of phage virion proteins with a large pretrained protein language model and an MLP neural network

**DOI:** 10.1101/2023.12.29.573676

**Authors:** Bowen Li, Guanxiang Liang

## Abstract

Bacteriophages, also known as phages, are essential for the stability of the microbiome system due to their ability to infect prokaryotes, another significant component of the microbiome. Thus, understanding the functions of phage proteins could help us unravel the nature of phages and their roles in the microbiome. However, limited by the low throughput of experimental techniques, a vast number of phage proteins remain unannotated in terms of their functions. Computational methods are expected to solve this restriction due to their high throughput and cost-effectiveness. In this study, we focused on one aspect of functional annotation for phage proteins, the identification and classification of phage virion proteins, and the integration of a large pretrained protein language model and an MLP neural network dramatically improved the performance of these two tasks. Additionally, we compared our model with some previous deep learning models using a newly collected, independent benchmark dataset, demonstrating the strong generalization ability of our model for both tasks. The source codes of ESM-PVP and the software for the PVP identification task have been uploaded to: https://github.com/li-bw18/ESM-PVP.

## 1. Introduction

With the large-scale application of metagenomic or virome sequencing and the pipelines for genome assembling, a large number of putative viral genomes have been assembled from samples of different origins (Camarillo-Guerrero et al. 2021; Roux et al. 2016). Of these assembled genomes, a significant portion of them are identified or predicted as phage genomes, and meanwhile, as phages are able to infect prokaryotes, they might be critical in maintaining the stability of the entire microbiome (Dion et al. 2020; Gregory et al. 2020; Salmond and Fineran 2015). In addition, phages have applications in both scientific research and daily life, such as synthetic biology, gene editing technology, agricultural control, sewage treatment, food safety, phage therapy, and so on (Fernández et al. 2018; Salmond and Fineran 2015). Therefore, deep exploration of the properties and functions of these phages is of great significance for us to understand the role of phages in the ecosystem and better apply them to the above fields. To achieve this, one of the initial steps is to annotate the functions of proteins from these assembled phage genomes. However, traditional experimental techniques face challenges in keeping pace with the rapid generation of new data. Moreover, functional annotation based on sequence alignments is not suitable for a large proportion of completely novel proteins, resulting in the emergence of a large number of proteins awaiting annotation, also known as functional dark matters (Fitzgerald et al. 2021; Liang and Bushman 2021; Pavlopoulos et al. 2023). Independent or weakly dependent on the sequence similarity, machine learning, and deep learning methods have the potential to annotate these functional dark matters, and at the same time, they have a very high throughput when the computing power is sufficient and could predict the functions for a large number of proteins in an acceptable time and with a less cost.

An important task for the functional annotation of phage proteins is the identification and classification of phage virion proteins (PVPs), also called phage structural proteins. The outer shell of a phage, which is used to protect its genetic material, is made up of phage virion proteins, and the mechanisms of how phages recognize their hosts are also implicit in the functions of these proteins (Kabir et al. 2022). Therefore, in addition to the significance mentioned above, annotation of PVPs is also valuable for discovering how phages infect their hosts, predicting the hosts for phages, and developing antimicrobial drugs based on phage proteins (Shang et al. 2023).

There are several machine learning and deep learning methods for the identification and classification of PVPs. Traditional machine learning models utilize different feature extraction methods with various classifiers, such as Random Forest (RF), Naive Bayes (NB), Support Vector Machine (SVM), Gradient Boosting Classifier (GBC), Scoring Card Method (SCM), Light Gradient Boosting Machine (LGBM) and so on. Among these methods, most only utilize one single classifier: RF_phage virion (Zhang and Li 2023) and the model from Ru et al. (2019) adopt the RF classifier; PhagePred (Pan et al. 2018) and the model from Feng et al. (2013) utilize the NB probabilistic model; the model from Zou and Yu (2023), Pred-BVP-Unb (Arif et al. 2020), PVP-SVM (Manavalan et al. 2018), the model from Tan et al. (2018), and PVPred (Ding et al. 2014) use the SVM framework; the model from Barman et al. (2023) applies the GBC; Phage_UniR_LGBM (Bao et al. 2022) employs the LGBM; PVPred-SCM (Charoenkwan et al. 2020a) leverages the SCM. Others contain more than one classifier and exploit the ensemble methods to improve the performance, including SCORPION (Ahmad et al. 2022), iPVP-MCV (Han et al. 2021), Meta-iPVP (Charoenkwan et al. 2020b), and the model from Zhang et al. (2015).

The first deep learning method for PVP identification is iVIREONS (Seguritan et al. 2012), which utilizes the Multi-Layer Perceptron (MLP) with the features of amino acid frequency, however, it could not classify PVPs into detailed classes (the classification task). Another method without the classification task is VirionFinder (Fang and Ding 2021), which leverages the Convolutional Neural Network (CNN) with both the protein sequence feature and the biochemical property feature to accomplish the identification task. PhANNs (Cantu et al. 2021) is the very first method containing the classification task, which exploits the MLP neural network and multiple features related to the sequences and the chemical properties to identify PVPs and additionally classify them into ten classes. DeePVP (Fang et al. 2022) uses the same training dataset as PhANNs, performs the same tasks and innovates in the model, and with the strategy of the natural language processing (NLP), DeePVP uses the CNN model to make predictions completely from the sequences. PhaVIP (Shang et al. 2023), instead, creatively introduces the technology of image processing and the latest Vision Transformer (ViT) architecture to these tasks, and in detail, it applies the chaos game representation (CGR) to encode protein sequences into images, and utilizes the ViT which has recently shone in image tasks for the final prediction.

Recently, Transformer-based language models have been widely used in the NLP, and problems related to the biological macro-molecule could also be applied to the paradigm of NLP, so a large number of protein or nucleic acid language models have been developed to solve a variety of biological problems (Ji et al. 2021; Lin et al. 2023). When many layers of Transformer are included in a single language model, it is not possible to train such a model from scratch due to limitations of the current training technology, so in this case, pretraining technology is necessary. ESM (Lin et al. 2023; Rivers et al. 2021), developed by Meta AI, is a large pretrained protein language model based on the BERT framework, and combined with a simple classifier, various protein-related downstream problems could be completed by finetuning or using the embedding of the pretrained ESM model, for example, NeuroPred-PLM (Wang et al. 2023) finetunes the ESM-1 with 85M parameters (a version of ESM) and adopts a classifier which consists of both the CNN and attention layers to improve the performance for the prediction of neuropeptides; CLEAN (Yu et al. 2023) uses the embedding of the ESM-1b with 650M parameters and constructs a classifier training by contrastive learning to accurately predict the EC numbers of enzymes.

Here, we built ESM-PVP with the finetuned ESM-2 (650M parameters) and an MLP neural network to both identify and classify PVPs. The training process directly utilized the dataset from PhaVIP, which is the latest PVP dataset with reasonable processes of data filtering and splitting (Shang et al. 2023). ESM-PVP just accepts a protein sequence as the input, and the sequence is processed into integer tokens, and then ESM-2 module converts these tokens into the sequence embedding, which is fed into the MLP network to obtain the final prediction. In line with the usual practice of finetuning processes, during training, some of the pretrained parameters in the ESM-2 module were finetuned, while the others remained the same, and the parameters in the MLP were all trained from scratch. The structures of the MLP and the details of training are slightly different in the identification (binary) and classification (multi-class) tasks for better adaptation to specific issues. In addition to the standard test, we collected an independent test set from RefSeq to benchmark most previous deep learning models and our model as well, and all of the sequences in this set were released in 2023, that is, none of these models had utilized these sequences for training, further comparing the robustness of these methods. Overall, ESM-PVP demonstrates better performance as well as stronger generalization capabilities, illustrating the incredible power of large protein language models.

## 2. Materials and Methods

### 2.1 The dataset for model training

The PhaVIP dataset was collected from NCBI RefSeq in 2022 utilizing strict selection criteria to guarantee the data quality, and finally, it contains more than 80,000 phage protein sequences, with almost balanced positive (PVPs) and negative (non-PVPs) samples, and for the detailed classification of PVPs, there are eight different categories: the baseplate, major capsid, major tail, minor capsid, minor tail, portal, tail fiber, and other (Shang et al. 2023). PhaVIP split this dataset into training and test sets by two different strategies: one was the time split, which grouped the data released before December 2020 into the training set, and after this time into the test set; another was the cluster split, which divided the dataset into two sets according to different similarity thresholds, making the sequences in the two sets less similar (Shang et al. 2023). Models trained with the time split sets tend to have a stronger generalization ability, while those trained with the cluster split sets are better at predicting sequences with limited homologous proteins in the references. The authors of PhaVIP published the time split sets used in the training of the binary task and the cluster split sets applied in the training of the multi-class task. In order to compare more convincingly with the previous methods, we just leveraged the above two kinds of sets to complete the training of binary and multi-class tasks, respectively. It should be noted here that the samples of the other class were missing from the multi-class sets we obtained, so our multi-class task could only classify seven categories, but we took this bias into consideration and tried to dispose of it carefully when comparing with other models (later mentioned). Finally, for the binary task, there are 27,704 PVPs and 36,778 non-PVPs in the training set, while 7,509 PVPs and 10,103 non-PVPs in the test set, and for the multi-class task, there are six groups of training and test sets with the similarity thresholds of 0.4, 0.5, 0.6, 0.7, 0.8, and 0.9.

### 2.2 Model structures and training processes

The structures of ESM-PVP are shown in **Fig. 1**. For the binary part of this model, at first, a phage protein sequence, which has been converted into tokens, enters into the ESM-2 with 33 Transformer Encoder layers and 650M parameters, and is encoded into embedding, which contains a wealth of information about the properties of the protein (Lin et al. 2023). Then the embedding sums in the dimension of the sequence, forming a vector with a length of 1280, and this vector enters into an MLP with 4 layers, which compresses the vector and extracts the useful information. Finally, the probability value of PVP is obtained through the Softmax activation function, and in general, when the probability value is greater than 0.5, the protein is considered to be a PVP. Some ReLU activation functions are also added into the model for improving the nonlinear expression ability and the performance of the model. During training, the ESM-2 module loaded the pretrained parameters from Meta AI, and except for the last four layers of Transformer Encoder, the parameters of the other layers were frozen and could not be adjusted, while the parameters in the MLP were trained from the initialization state. The learning rate was set to 5e-6, the batch size was 2, and a weight decay of 1e-5 was also applied. 20% of the training sets were utilized to valid the model, and with only 1 epoch, the model converged.

**Figure 1.**
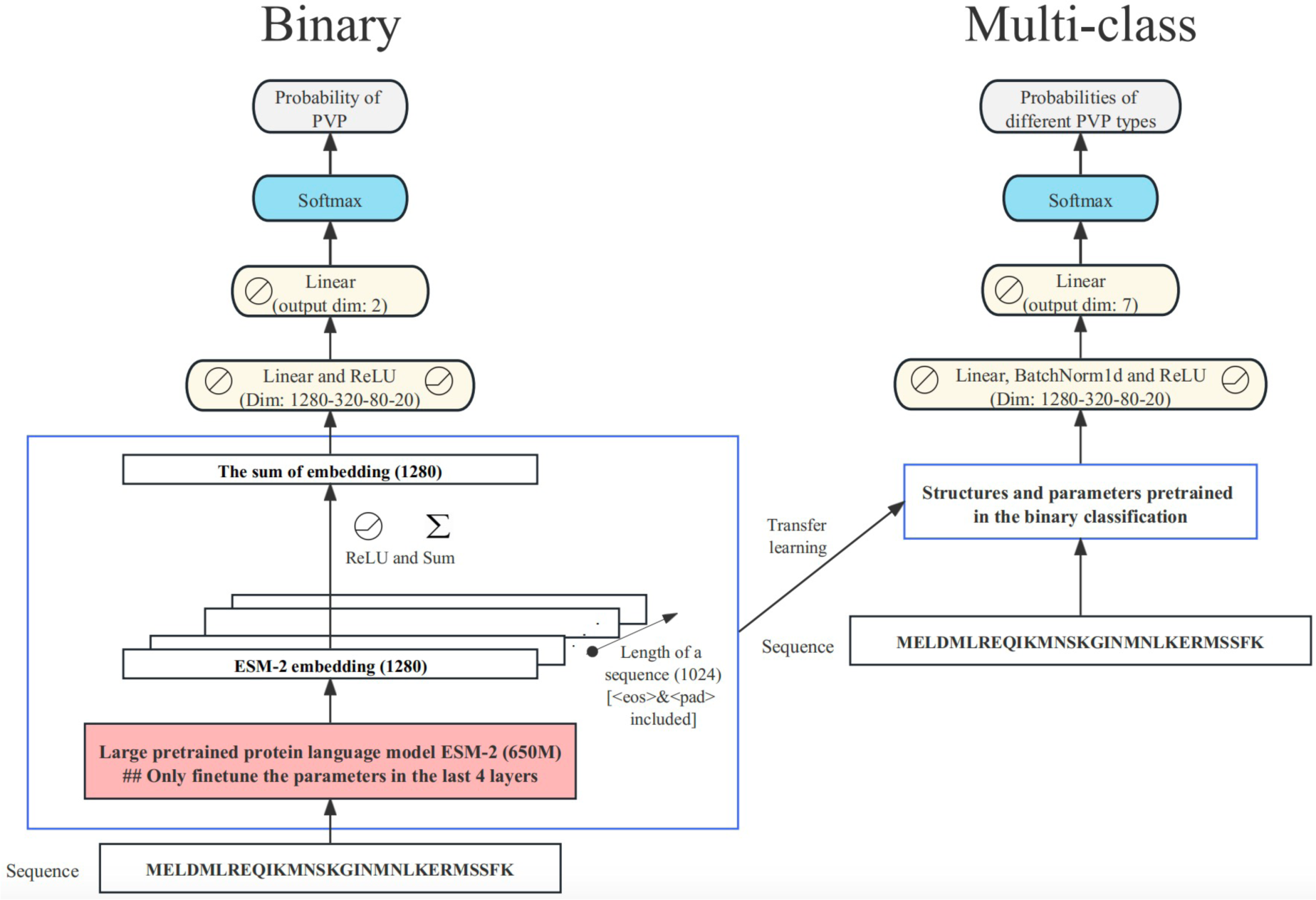
The structures of ESM-PVP. The left panel shows the binary (identification of PVPs) part, and the right panel shows the multi-class (classification of PVPs) part. The dimensional changes during the model operations are indicated in each module. When a protein sequence is imported, some special characters, such as the end character <eos> and the completion character <pad> might be added into the sequence, and the excessively long sequence might be truncated, and the final input sequence length is uniformly 1024.

For the multi-class part of this model, due to the large number of categories, it is necessary to choose a large batch size, but this is a huge challenge when computing resources are limited. To balance the computing power and the performance, we exploited the transfer learning strategy, and the parameters in the ESM-2 of the multi-class part directly utilized the parameters trained in the binary part. In practice, we completely separated the ESM-2 from the following MLP classifier. The sequence is first inferred in the binary part, and the intermediate vector before the binary MLP is taken out as the input of a separate multi-class MLP classifier. This effectively reduced the memory consumption during training, because only the multi-class MLP was involved in the forward propagation, backpropagation, and updates of parameters. We added some 1D Batch Normalization layers to the multi-class MLP to train the model more stably and improve the performance in the case of a larger batch size. The learning rate was set to 1e-4, the batch size was 64, a weight decay of 1e-5 was applied, and the weighted loss was used for the imbalance of samples from different classes. For each sequence similarity threshold, an independent model was trained with 20% of the training sets randomly selected for validation, and these models could converge after about 30 epochs.

### 2.3 Test processes and metrics

We ran two sets of tests on the model, and the typical test utilized the test set separated from PhaVIP dataset (mentioned before), while another independent test used a test set newly collected by us. In detail, we obtained all PVPs released in 2023 (2023/1/1-2023/12/15) from NCBI RefSeq, randomly selected 48 proteins for each class in the multi-class task, in total 336 PVPs, and meanwhile, collected 336 non-PVPs in a similar way as negative samples for the binary task. In this way, we each constructed a balanced test set for the binary and multi-class tasks. Since PhaVIP authors had already fully tested other deep learning methods with the first test set, we only additionally included the PhaVIP model during the first test (Shang et al. 2023). The second test included all the deep learning models that we were confident to run correctly to compare the robustness of different models.

A variety of metrics were chosen to evaluate and compare the performance of models. For the binary task, we might calculate the accuracy (ACC), precision, recall, F1, AUC, and AP value. The formulas for ACC, precision, recall, and F1 are as follows (1)-(4):

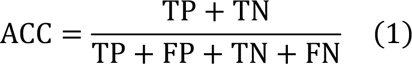

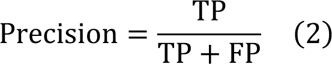

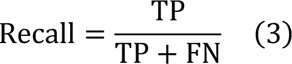

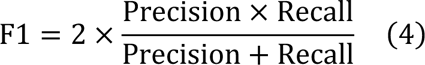

where TP, FP, TN, and FN mean the number of true positive, false positive, true negative, and false negative samples during a test. The above metrics are typically calculated under the premise that the probability threshold is 0.5, but when the threshold of classification is adjusted, totally different values of these metrics will be obtained. With different values under different thresholds, the ROC and PR curve could be plotted, and the ROC curve indicates the relationship between the true positive rate (TPR) and false positive rate (FPR), while the PR curve indicates the relationship between the precision and recall. The formulas for TPR and FPR are as follows (5)-(6).

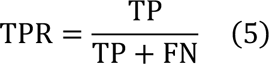

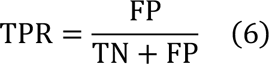

The AUC represents the area under the ROC curve, and the AP represents the area under the PR curve. Combining the above metrics could evaluate and analyze the performance of different models from multiple perspectives, especially the F1, AUC and AP metrics.

For the multi-class task, when a certain class is treated as positive, while other classes are all treated as negative, the precision, recall, and F1 could also be calculated for this class. Therefore, when we tested the multi-class task, we might calculate the average precision, recall, and F1 (also called macro-F1) over different classes, and ACC (also called micro-F1) was also considered.

### 2.4 Computing resources

Up to eight 40G NVIDIA A40 and two 32G NVIDIA Tesla V100 PCle GPUs were utilized for model training and inference, and these GPUs were all from the public platform of School of Medicine, Tsinghua University.

## 3. Results

The performance of the binary part of ESM-PVP on the PhaVIP test set is shown in **Fig. 2** and **Table. 1**. Due to the possible differences in the device and validation set selection, we retrained the PhaVIP model to eliminate the effects of the above problems, and the performance of retrained PhaVIP is also shown in **Fig. 2** and **Table. 1**. It could be seen that ESM-PVP performs slightly better than PhaVIP in all metrics.

**Figure 2.**
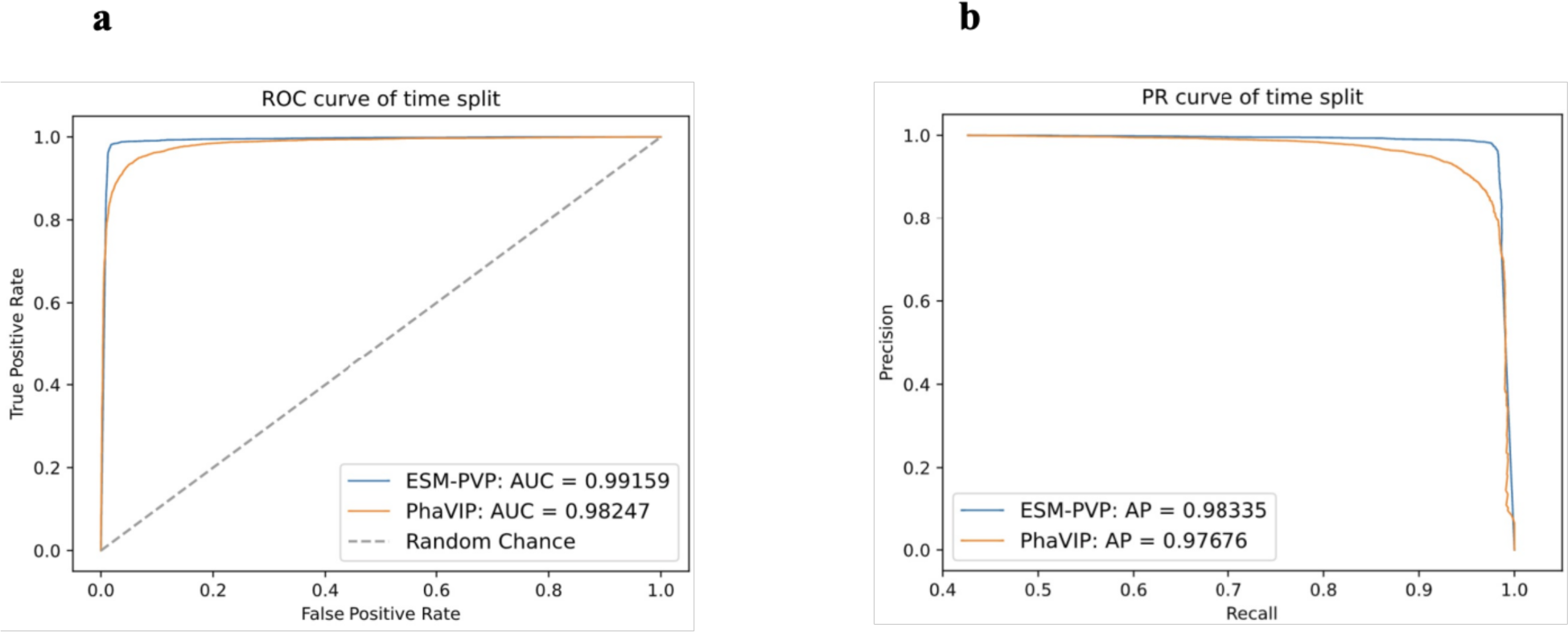
The performance comparison of ESM-PVP and PhaVIP on the identification task with the PhaVIP test set. a, the ROC curve of two methods; b, the PR curve of two methods. The AUC and AP scores are annotated in the figure.

**Table 1.** The precision, recall, and F1 of ESM-PVP and PhaVIP on the identification task with the PhaVIP test set.

| **Metric** | Precision | Recall | F1 |
| --- | --- | --- | --- |
| ESM-PVP | 0.9770 | 0.9778 | 0.9774 |
| PhaVIP | 0.9308 | 0.9344 | 0.9326 |

The performance of the multi-class part of ESM-PVP on the PhaVIP test set is shown in **Fig. 3**. For the absence of the other class, we were unable to retrain the multi-class part of PhaVIP, so during the comparison, the performance of PhaVIP was directly obtained from that reported by the authors of PhaVIP, and we could discover that ESM-PVP outperforms the PhaVIP in the macro-F1 score at every sequence similarity threshold (**Fig. 3a**). More interestingly, ESM-PVP does not seem to depend on the sequence similarity at all, with little difference in the performance of different thresholds, suggesting the powerful capabilities of large protein language models for the functional prediction of proteins with none or few homologous proteins (**Fig. 3a**). As for the single class, the performances of different classes vary, and the minor capsid class, which has the smallest sample number, performs the worst among seven classes, indicating that ESM-PVP is still unable to cope with classes with insufficient samples (**Fig. 3b**).

**Figure 3.**
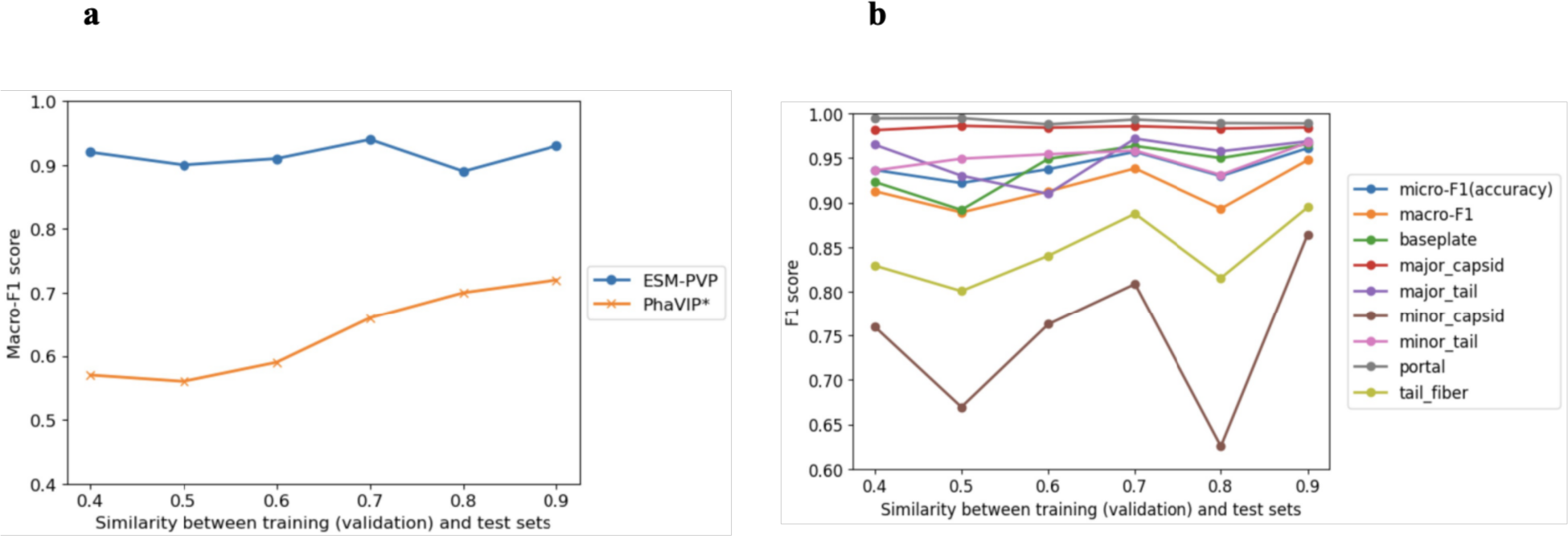
The performance comparison of ESM-PVP and PhaVIP on the classification task with the PhaVIP test set. a, the macro-F1 score of two methods at different sequence similarity thresholds; b, the F1 score of each class in the ESM-PVP. *The metrics of PhaVIP were obtained from the supplementary materials of PhaVIP.

The benchmark of an independent test set is meaningful for assessing the robustness of different models, and in this section, we tested all the deep learning models that had been explicitly packaged into software or a webserver. Note that this limitation is because our experience has shown that it is often difficult to run correctly with models that are not well packaged, failing to reflect the real performances. The results are shown in **Fig. 4**. The PhANNs and PhaVIP predicted these proteins on their webservers, and DeePVP on a virtual machine provided by the authors. In the binary task, except for the precision value of DeePVP, ESM-PVP performs the best in other metrics (**Fig. 4a**). In the multi-class task, there were ten classes in the outputs of DeePVP and PhANNs, including all the seven classes we used and another three classes, as a result, to eliminate this bias, for those samples with predictions of the other three classes, we adjusted their predictions to the one with the highest probability value in our seven classes for the calculation of metrics. As for PhaVIP, there was an extra other class in its outputs, but since the probability values of all classes were not provided, for those samples predicted to be the other class in PhaVIP, we could not obtain the class with the second highest probability value as a substitute, so for these samples, we directly assumed that they were correctly predicted, which ensured that the effect of PhaVIP was not underestimated. **Fig. 4b** demonstrates that ESM-PVP performs the best across all metrics. Overall, ESM-PVP shows excellent robustness during the benchmark, suggesting that it is capable of making predictions on the newly discovered phage proteins.

**Figure 4.**
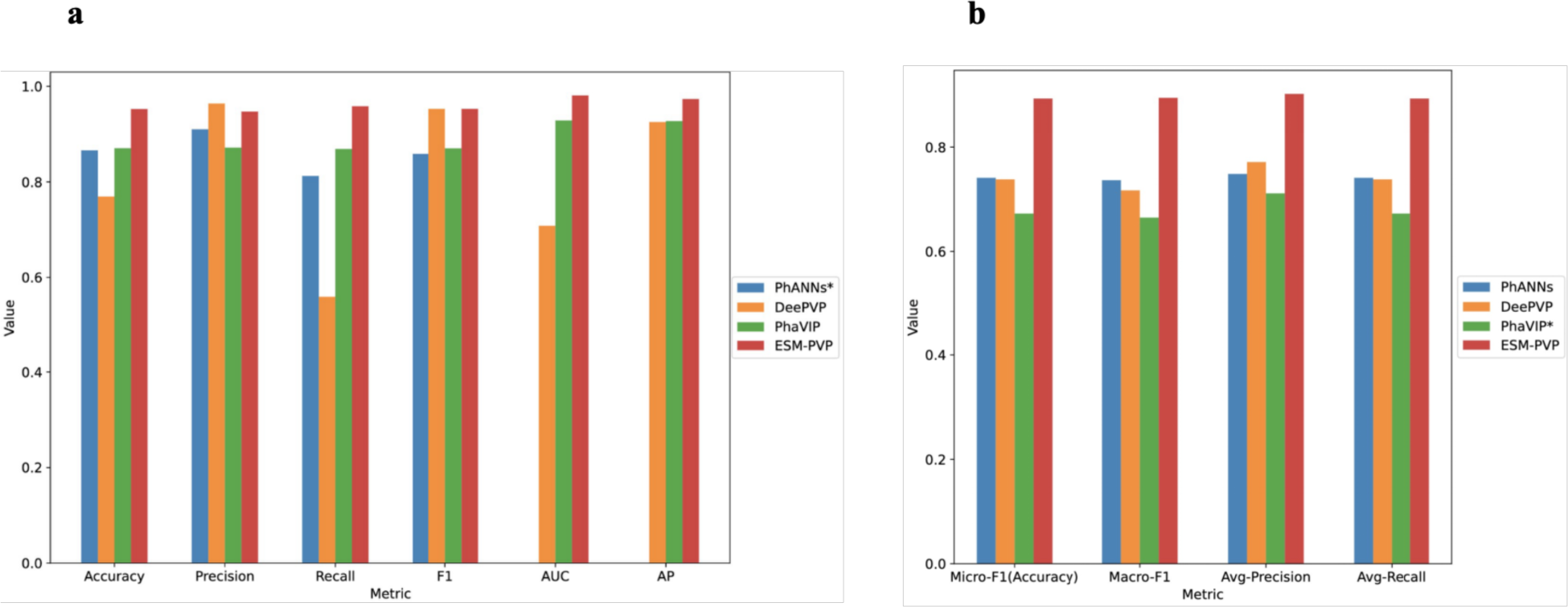
The performance comparison of some deep learning models with the independent test set. a, the metrics of the binary task; b, the metrics of the multi-class task. *The outputs of PhANNs webserver do not include the probability of PVP or non-PVP, so the AUC and AP values of PhANNs cannot be calculated in the left panel; the outputs of PhaVIP webserver do not include the probabilities of all classes, so for those proteins annotated as other class by PhaVIP in the multi-class task, they are directly considered to be correctly predicted, which might overestimate the performance of PhaVIP.

## 4. Discussions

Although ESM-PVP performs well on two tasks, there is still room for improvement, especially for the multi-class task. To determine the shortcomings of ESM-PVP, we calculated the confusion matrix of ESM-PVP on the independent test set and plotted heatmaps (**Fig. 5**). In the binary task, there is no significant difference in the number of false positive and false negative samples, proving that ESM-PVP does not have a significant bias (**Fig. 5a**). In the multi-class task, most errors come from the minor capsid proteins, which results from the insufficient number of training samples for the minor capsid class (**Fig. 5b**). This suggests that how to improve the performance of these classes with insufficient data number, also including those classes that were discarded due to too little data in PhaVIP (e.g. collar), is an important direction in the future.

**Figure 5.**
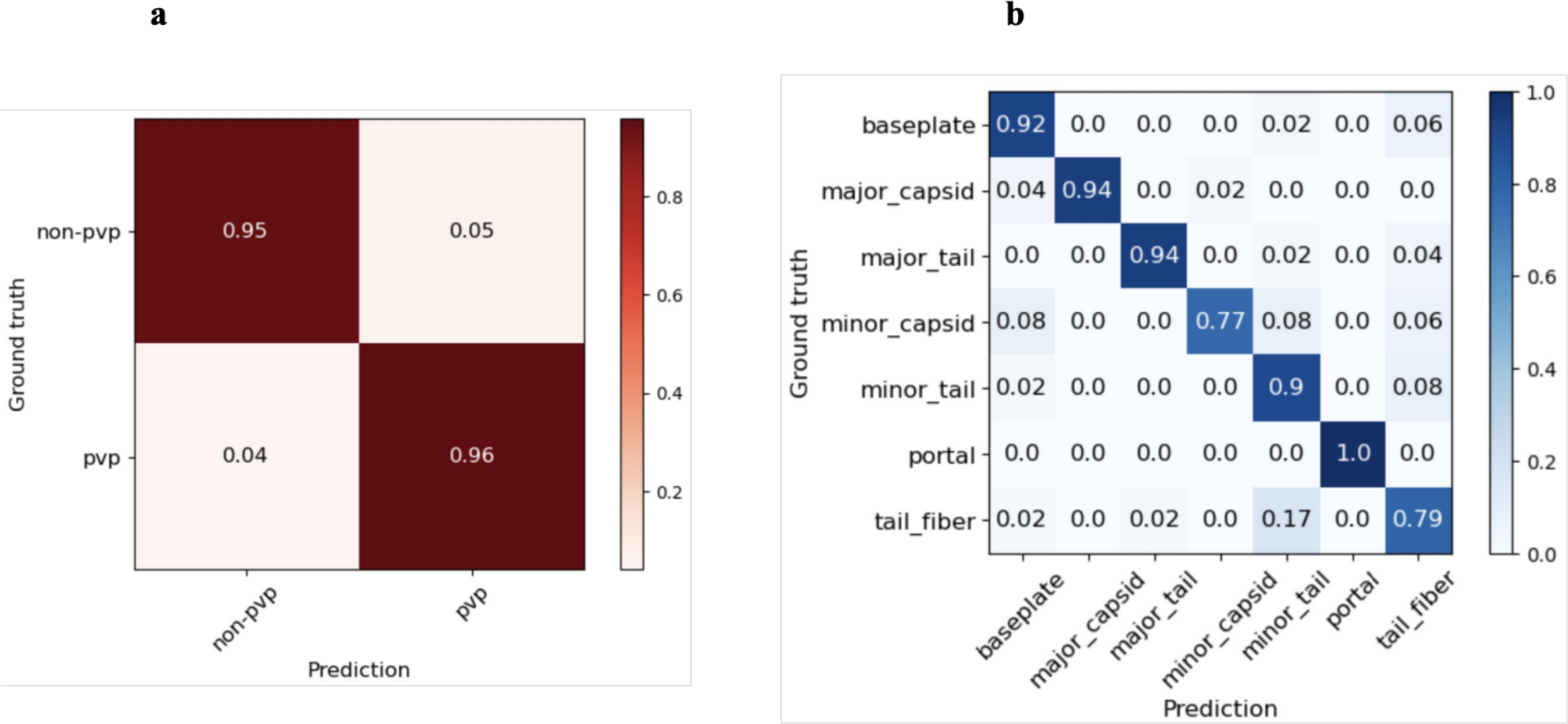
The confusion map of ESM-PVP on the independent test set. a, the binary task; b, the multi-class task.

Additionally, there are a large number of samples from other classes that are misclassified as the tail fiber class, and after simple analysis, we found that this might stem from the mislabeling of tail fiber proteins in the training data (**Fig. 5b**). Since when we were collecting the independent test set, we found some proteins in the tail fiber dataset that should be labeled as the minor tail (e.g. YP_010659469.1) class, as well as some proteins in the major tail or minor tail dataset that should be labeled as the tail fiber (e.g. YP_010747610.1, YP_010747626.1, YP_010674637.1) class, even when we had carefully obtained this test set by referring to the data collection methods of PhaVIP, PhANNs, and some recommendations from a review (Cantu et al. 2021; Kabir et al. 2022; Shang et al. 2023). Therefore, in order not to confuse the model, there might still be room for optimization in the data collection of PVP and non-PVP.

Finally, because of the absence for the other class in multi-class training set, the multi-class part of ESM-PVP could not be packaged into software now (the class is incomplete), so next we will collect a new dataset soon and retrain the multi-class part to make up for this lack.

